# In vivo growth of *Staphylococcus lugdunensis* is facilitated by the concerted function of heme and non-heme iron acquisition mechanisms

**DOI:** 10.1101/2022.01.12.476141

**Authors:** Ronald S. Flannagan, Jeremy R. Brozyna, Brijesh Kumar, Lea A. Adolf, Jeffrey John Power, Simon Heilbronner, David E. Heinrichs

## Abstract

Acquisition of iron underpins the ability of pathogens to cause disease and *Staphylococcus lugdunensis* has increasingly been recognized as a pathogen that can cause serious infection. In this study, we sought to address the knowledge gap that exists regarding the iron acquisition mechanisms employed by *S. lugdunensis*, especially during infection of the mammalian host. Here we show that *S. lugdunensis* utilizes diverse genome encoded iron acquisition mechanisms to satisfy its need for this nutrient. Indeed, *S. lugdunensis* can usurp hydroxamate siderophores, and staphyloferrin A and B from *S. aureus*, using the *fhuC* ATPase-encoding gene. Acquisition of catechol siderophores and catecholamine stress hormones necessitates the presence of the *sst-1* transporter-encoding locus, but not the *sst-2* locus. Iron-dependent growth in acidic culture conditions necessitates the *feoAB* locus. Heme iron is acquired via expression of the iron-regulated surface determinant (*isd*) locus. During systemic infection of mice we demonstrate that while *S. lugdunensis* does not cause overt illness, it does colonize and proliferate to high numbers in the kidneys. By combining mutations in the various iron acquisition loci, we further demonstrate that only a strain mutated for all of *isd*, *fhuC, sst-1,* and *feo*, versus combination mutants carrying wild type copies of any one of those loci, was attenuated in its ability to proliferate to high numbers in kidneys. Taken together our data reveal that *S. lugdunensis* requires a repertoire of both heme and non-heme iron acquisition mechanisms to proliferate during systemic infection of mammals.

**Importance:** Acquisition of iron underpins the ability of pathogens to cause disease and *Staphylococcus lugdunensis* has increasingly been recognized as a pathogen that can cause serious infection. In this study, we sought to address the knowledge gap that exists regarding the iron acquisition mechanisms employed by *S. lugdunensis*, especially during infection of the mammalian host. Owing to an inability to synthesize siderophores, growth of *S. lugdunensis* is dramatically impaired in the presence of transferrin or serum, yet S*. lugdunensis* nonetheless uses several other genome-encoded iron acquisition mechanisms, in concert, to proliferate within the mammalian host. Therefore, the development of interventions that target bacterial iron acquisition systems should consider the overlapping function of distinct metal acquisition strategies deployed by bacterial pathogens.

## Introduction

Iron (Fe) is an essential nutrient for nearly all forms of life and despite its importance and abundance on earth it primarily exists, at neutral pH, in an insoluble, ferric iron (Fe^3+^) state. In the context of the host, free Fe is scarce and is maintained at a concentration well below the requirement needed to support microbial growth (1). Within the host most Fe is contained within heme prosthetic groups in hemoglobin inside circulating erythrocytes (2). Fe may also be sequestered within host cells by the iron-storage protein ferritin or bound by extracellular serum glycoproteins such as transferrin and lactoferrin (3). Collectively these iron binding proteins sequester Fe to minimize toxicity and to purpose this trace metal for host cellular processes. In addition, these proteins along with other immune effectors including hepcidin (4, 5), ferroportin (4, 6), and calprotectin (7, 8), collectively act to limit Fe availability to invading microorganisms through a process termed nutritional immunity (9, 10). Therefore, to successfully colonize and infect a host, an invading bacterial pathogen must overcome host-driven nutrient sequestration and acquire Fe to support growth *in vivo* (1).

*Staphylococcus lugdunensis*, like several other coagulase-negative staphylococci (CoNS), is a human skin commensal and even protect against colonization by *S. aureus* (11). However, it can also act as a pathogen that displays elevated virulence as compared to other CoNS (12–15). Indeed, infections caused by *S. lugdunensis* are reminiscent of those attributed to *S. aureus* and, in a susceptible host, *S. lugdunensis* can cause a spectrum of infections including skin and soft tissue infections (SSTIs), bacteremia, pneumonia, and osteomyelitis (12, 13, 16, 17). In addition, *S. lugdunensis* has a propensity to cause aggressive infective endocarditis with mortality rates that can be as high as 50% (18, 19). The ability of *S. lugdunensis* to cause infection requires the concerted action of numerous virulence factors (20, 21) and necessitates that this bacterium acquires Fe from its host.

Successful pathogens can deploy a variety of mechanisms to acquire Fe from the host and this can involve extracting Fe from both heme and non-heme Fe sources to support growth during infection (22, 23). Indeed, in Gram-positive bacteria such as *S. aureus*, the iron-regulated <u>s</u>urface <u>d</u>eterminant (Isd) system functions to shuttle heme from the extracellular milieu across the bacterial cell wall and cytoplasmic membrane where Fe atoms can be extracted (23–28). *S. lugdunensis* is unique among CoNS in that it also encodes a functional Isd system (29–32) and utilizes this high affinity heme uptake pathway to grow at low (<500 nM) heme concentrations (33–36). In addition, *S. lugdunensis* utilizes a high affinity energy coupling factor (ECF) type transporter named Lha to extract heme from diverse host hemoproteins (37). To acquire non-heme Fe, most bacteria produce low molecular weight high affinity iron chelators termed siderophores (38). Through siderophore production bacteria can extract Fe^3+^ from oxyhydroxide precipitates or, for pathogens, from host glycoproteins such as transferrin; siderophore production has been shown in many bacteria including *S. aureus* to contribute significantly to pathogenesis *in vivo* (38). *S. aureus* elaborates two carboxylate-type siderophores, staphyloferrin A (SA) and staphyloferrin B (SB), of which the biosynthetic proteins are encoded by *sfa* and *sbn* loci, respectively (39–41) SA and SB are transported by the ABC transporters HtsABC and SirABC, respectively, which are encoded by loci adjacent to their cognate siderophore biosynthetic genes (42, 43). Contrary to *S. aureus*, *S. lugdunensis* does not produce either SA or SB (44), however, *S. lugdunensis* expresses the transporters HtsABC and SirABC and can thus usurp SA and SB produced by *S. aureus* (44). *S. aureus* can also utilize xenosiderophores (i.e. siderophores produced by other microbes) and their utilization requires expression of the ferric <u>h</u>ydroxamate <u>u</u>ptake (Fhu) transporter and the Sst catechol transporter (45, 46). While Fhu enables *S. aureus* to utilize hydroxamate-type siderophores, Sst allows *S. aureus* to utilize siderophores containing catechol/catecholamine moieties (39, 47, 48). Host-derived stress hormones such as epinephrine and dopamine are catecholamines that chelate Fe and act as ‘pseudosiderophores’ to bacteria expressing catechol transport systems (46, 49–52). Indeed, *S. aureus* utilizes the Sst pathway for catechol utilization but Sst functionality is only evident when endogenous biosynthesis of SA and SB is perturbed (46).

In comparison to *S. aureus* there exists a paucity of information regarding the Fe acquisition mechanisms employed by *S. lugdunensis*, especially during infection of the mammalian host. The lack of information on *S. lugdunensis* takes on added significance when one considers the pathogenic potential of this microbe. To rectify this, we investigated the iron procurement strategies of *S. lugdunensis* both *in vivo* and *in vitro*. We demonstrate that *S. lugdunensis* encodes and utilizes the Fhu and Sst transport proteins to acquire iron from a variety of hydroxamate and catechol/catecholamine siderophores, and that the ferrous iron transport system, encoded by the *feoAB* genes, functions in *S. lugdunensis* to acquire iron under acidic culture conditions. During systemic infection of mice with *S. lugdunensis*, the bacteria seed the kidneys where they subsequently proliferate to high numbers. We demonstrate that growth of *S. lugdunensis* in the murine kidney requires the concerted action of both heme (i.e. Isd) and non-heme (including *feo*) iron acquisition systems.

## Results

### *S. lugdunensis* is restricted by serum *in vitro* yet the bacteria proliferate *in vivo*

Human infection by *S. lugdunensis* can be severe and this coagulase-negative *Staphylococcus* spp. is often erroneously identified as *S. aureus*. Given that the ability to acquire iron from the host is essential for bacteria to cause infection, we speculated that *S. lugdunensis* and *S. aureus* may display similar capacity for iron acquisition. Remarkably, these two related yet distinct bacterial species demonstrate profound differences with respect to growth under conditions of iron restriction *in vitro* (Fig. 1A). Indeed, comparison of *S. aureus* USA300 and *S. lugdunensis* HKU09-01 growth in RPMI supplemented with casamino acids (RPMI-CAS) and increasing amounts of horse serum (HS), a source of iron-chelating transferrin, revealed that *S. lugdunensis* failed to grow in as little as 0.5% (v/v) HS (Fig. 1A). The inability of *S. lugdunensis* to grow in the presence of HS was rescued upon supplementation of the culture medium with 20 μM FeCl_3_ establishing that the growth defect was due to iron restriction (Fig. 1A). In contrast, *S. aureus* USA300 grew robustly in 40x as much HS indicating these two bacterial species display profound differences in their ability to acquire iron *in vitro*. Despite this discrepancy, *S. lugdunensis* can cause serious infection and therefore we posited this bacterium must employ mechanisms of iron acquisition that permit growth *in vivo*. To test this notion, we next performed systemic murine infection experiments where the ability of *S. lugdunensis* HKU09-01 to colonize and grow in the kidneys and liver of infected animals was evaluated up to 8 days post-infection (Fig. 1B and C). The murine kidney and liver were selected for this analysis as *S. lugdunensis* poorly infects the heart and spleen (Fig. S1). These experiments revealed that in the murine liver the burden of *S. lugdunensis* decreased over time and by day 4 post-infection an approximate 2-log reduction in bacterial burden was apparent (Fig 1B). In contrast, in the murine kidney it was evident that the burden of *S. lugdunensis* steadily increased over the 8-day infection where the bacterial counts increased by more than 2-log (Fig. 1C). These data indicate that *S. lugdunensis* HKU09-01 displays organ specific differences in bacterial proliferation and indicate that within the kidney *S. lugdunensis* must acquire iron within the murine host to support the significant bacterial growth.

**Figure 1.**
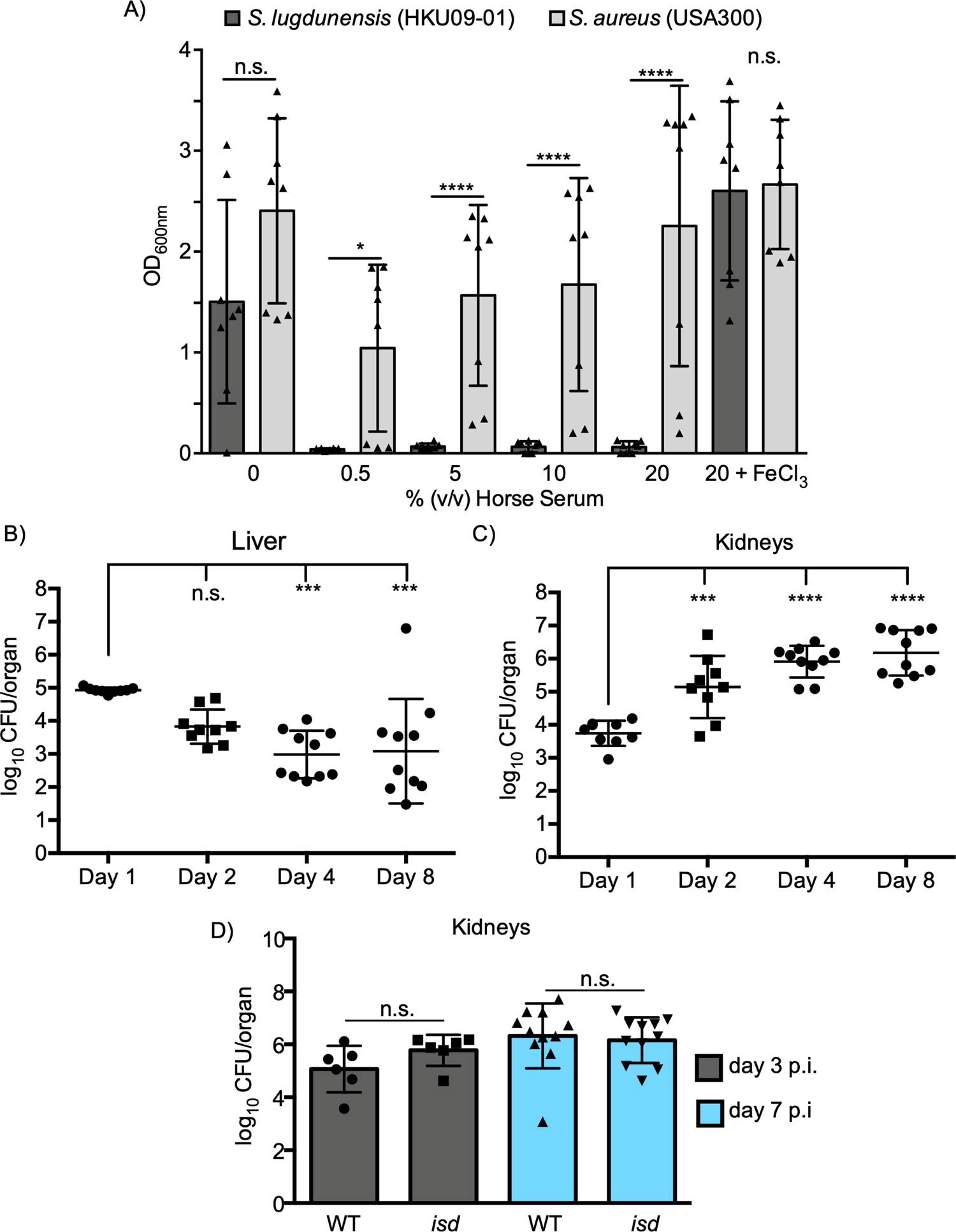
*S. lugdunensis* proliferates *in vivo* despite being restricted by serum. In (A), the growth of wildtype *S. lugdunensis* HKU09-01 and wildtype *S. aureus* USA300 in the presence of varying amounts of heat-inactivated horse serum in RPMI + 1% (w/v) casamino acids is shown. The data presented are the mean ± standard deviation of the endpoint optical density at 600nnm (OD600nm) measured after 24 hr. The data derive from three independent experiments and each symbol represents a separate biological replicate. Statistical significance was determined by performing an ordinary 2-way ANOVA. In (B) and (C) the burden of *S. lugdunensis* HKU09-01 in the kidneys and liver, respectively, of infected mice at different days post-infection is shown. In (C) the burden of *S. lugdunensis* HKU09-01 and an *isd* mutant at day 3 and day 7 post-infection is shown. In (B), (C) and (D) the data are presented as the mean log10 CFU/organ ± standard deviation and each data point represents an infected animal. Statistical significance was measured by ordinary one-way ANOVA with a tukeys multiple comparison post-test. n.s. denotes not significant and * p<0.05, *** p<0.001, **** p<0.0001.

### Isd deficient *S. lugdunensis* HKU09-01 does not display growth defects within the murine kidney

That *S. lugdunensis* HKU09-01 grows within the murine kidney prompted us to investigate whether the high affinity heme acquisition pathway encoded by the *isd* genes contributed to this phenotype. In *S. lugdunensis* HKU09-01 the entire *isd* locus is tandemly duplicated and previous work characterizing Isd function in *S. lugdunensis* has demonstrated that an increased copy number of the *isd* locus enhances *S. lugdunensis* growth through heme utilization (44, 53). Moreover, it has been demonstrated that heme may be accessible to staphylococci within the murine kidney as compared to other organs which could ostensibly support *S. lugdunensis* growth (54). To evaluate the contribution of the *isd* genes to growth in the murine kidney, animals were infected with wild-type *S. lugdunensis* or an isogenic *isd* mutant where the tandemly duplicated *isd* loci have been deleted. Importantly, *in vitro* growth of this strain in the presence of hemin (ferric chloride heme) is impaired relative to WT (Fig. S2). In vivo, however, *isd* deficient *S. lugdunensis* did not display a reduced bacterial burden in the murine kidney, as compared to the wildtype, at day 3 or day 7 post-infection indicating additional iron acquisition systems must help to support growth of *S. lugdunensis in vivo*.

### *S. lugdunensis* utilizes the ferric hydroxamate uptake (*fhu*) genes to acquire iron *in vitro*

Given *isd* is dispensable for *S. lugdunensis* growth within the murine kidney we sought to identify the additional iron acquisition systems that operate in this bacterium. Searches of the available genome sequences of *S. lugdunensis* identified genes homologous to the *S. aureus fhuCBG* locus that, in *S. aureus*, are required for ferric-hydroxamate siderophore-dependent iron acquisition (45). The proteins encoded by the *fhuCBG* genes in *S. lugdunensis* share significant identity to *fhuCBG* in *S. aureus* (see Table S1) and previous work has demonstrated that *fhuC* deficient *S. aureus* is debilitated for growth in iron deplete laboratory media and *in vivo* (48). This prompted us to assess the importance of *fhuC* during iron-restricted growth of *S. lugdunensis*. Analysis of the genomic sequence surrounding the *fhuCBG* locus in *S. lugdunensis* revealed that a canonical Fur box lies upstream of the *fhuCBG* locus suggesting the ferric iron <u>u</u>ptake <u>r</u>epressor (Fur) protein and cellular iron regulate transcription of these genes (Fig. 2A). In agreement with this notion, qPCR analysis revealed that the *fhu* genes in *S. lugdunensis* are significantly downregulated in response to Fe supplementation of the growth medium (Fig. 2B). This suggests that in low-iron environments, *S. lugdunensis* may utilize hydroxamate siderophores, if present, as a source of iron. To test this, we created an in-frame *fhuC* deletion in *S. lugdunensis* HKU09-01 to assess the role of *fhuC* in bacterial growth in the presence of deferoxamine (DFO) as a sole source of iron (Fig. 2C). Indeed, DFO can function as a xenosiderophore for *S. aureus* and chelate iron from transferrin owing to its exceptionally high affinity for Fe (55). This analysis revealed that *S. lugdunensis* lacking *fhuC* failed to utilize DFO for growth in the presence of HS and this growth defect could be rescued by supplementation of the growth medium with FeCl_3_. Moreover, provision of *fhuC* encoded on a plasmid also restored DFO utilization to the *fhuC* mutant establishing the observed growth defect in *S. lugdunensis* was attributable to *fhuC* inactivation alone (Fig. 2C).

**Figure 2.**
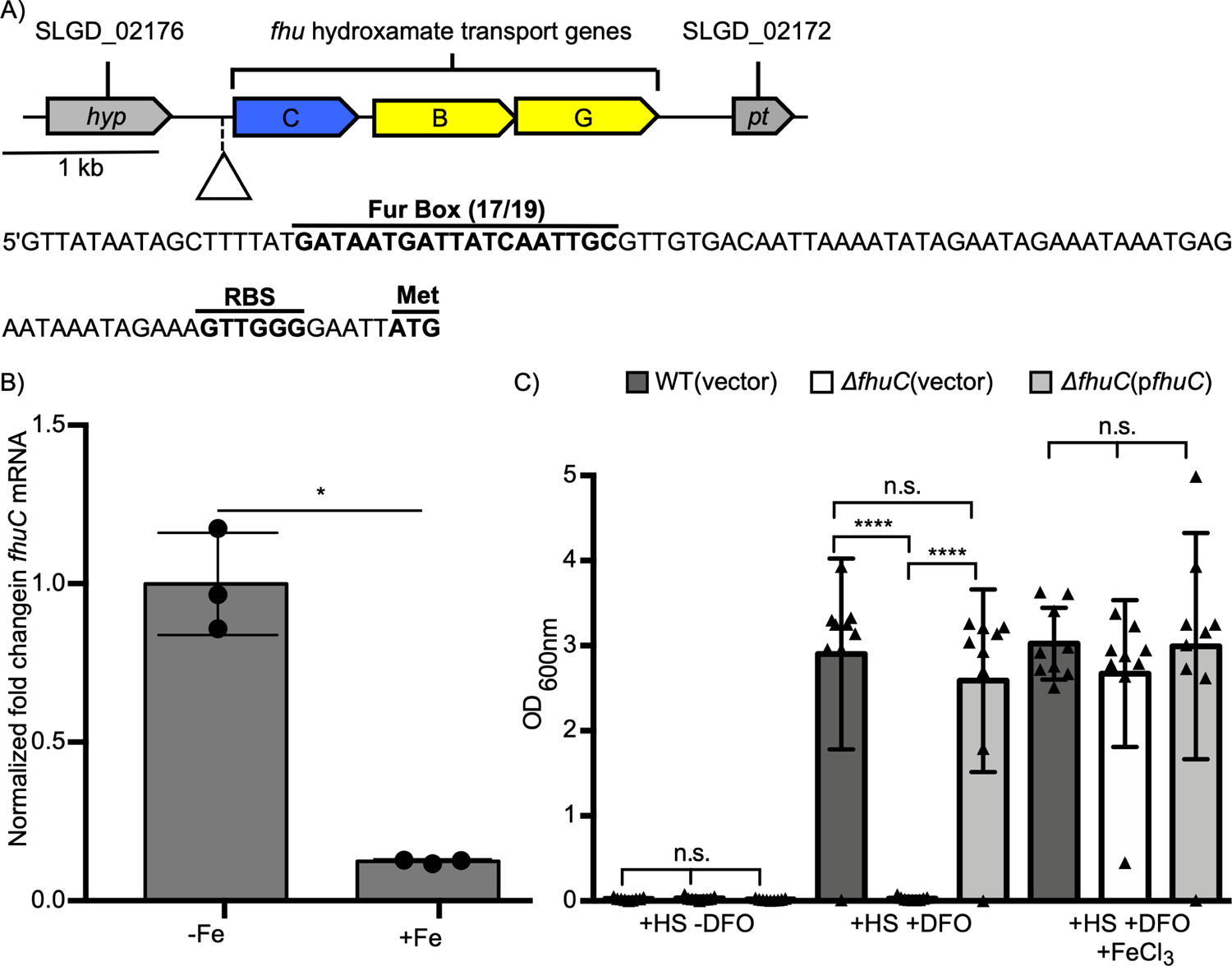
*Staphylococcus lugdunensis* deficient for th*e fhuC* gene fails to utilize deferoxamine (DFO) for growth under conditions of iron restriction. In (A) the physical map of the ferric-hydroxamate uptake genes in *S. lugdunensis* is shown. The promoter sequence for the operon is shown, with the putative Fur box, ribosome binding site (RBS) and start codon labeled. In (B) qPCR analysis of *fhuC* gene expression by wildtype *S. lugdunensis* grown in C-TMS (-Fe) or C-TMS with 100 μM FeCl_3_ (+Fe) is shown. Data was normalized relative to *rpoB* expression, and gene expression was normalized relative to that in C-TMS without added iron. The data derive from three independent experiments and a total of three biological replicates and statistical significance was determined by students t-test. In (C) the growth of wildtype *S. lugdunensis* HKU09-01 and Δ*fhuC* with either the vector control or the p*fhuC* plasmid. Growth was in RPMI + 1% (w/v) casamino acids with 0.05% (v/v) horse serum (HS) in the presence and absence of 100 μM DFO with or without 20 μM FeCl_3_. The data presented are the mean ± standard deviation of the endpoint optical density at 600nnm (OD600nm) measured after 24 hr. The data derive from three independent experiments and each symbol represents a separate biological replicate. Statistical significance was determined by ordinary one-way ANOVA with a tukey’s post-test. In (B) and (C) n.s. indicates not significant, *p<0.05, ****<p<0.0001.

In *S. aureus*, the FhuC protein functions as a promiscuous ATPase that provides energy for the transport of hydroxamate type xenosiderophores through the Fhu system, as well as the carboxylate siderophores, staphyloferrin A and B, that are transported into the cell through dedicated permeases encoded by the *htsABC* and *sirABC* operons, respectively (39, 42, 48, 56). In agreement with its predicted role in ferric hydroxamate uptake, the *fhuC* mutant was unable to use a variety of hydroxamate siderophores as an iron source in addition to DFO (Fig. S3). As expected, the *fhuC* mutant retained the ability to utilize catecholamine type siderophores and citrate (Fig. S3) that are transported through other dedicated siderophore uptake systems. Provision of the *fhuC* gene *in trans* eliminated the growth defects in the presence of hydroxamate siderophores confirming the importance of FhuC to xenosiderophore utilization. Of note, the *fhuC* mutant was also unable to utilize either SA or SB (i.e. carboxylate siderophores) when supplied as a sole source of iron (Fig. S3) indicating that, akin to *S. aureus*, *S. lugdunensis* utilizes the FhuC ATPase to energize uptake of SA and SB through HtsABC and SirABC, respectively (39, 48). Taken together these data reveal that *S. lugdunensis* HKU09-1 is reliant on the *fhuC* gene to utilize SA and SB, in addition to hydroxamate type siderophores, for growth under iron limited conditions.

Given the promiscuity of FhuC in effecting uptake of various siderophores, we wished to investigate whether it had any role to play in heme uptake. Within the *isd* locus is a gene encoding an ATPase (*isdL*). Strain HKU09-01 has a duplicated *isd* locus (>30 kb duplication) so to investigate this we used strain N920143 (single *isd* locus) to construct strains bearing mutations in *isdL* and *fhuC*. While the *isdL* mutant had a significant growth defect on hemoglobin as a sole iron source, the *fhuC* mutant did not, nor did it have any additive effect to *isdL* mutation on growth on hemoglobin as an iron source (Fig. S4A). Moreover, using yeast two-hybrid analyses, we demonstrated that IsdL, and not FhuC, interacted with the IsdF membrane permease (Fig. S4B), providing further proof that the Isd heme acquisition system has a dedicated ATPase to power heme uptake, and that FhuC does not function in heme acquisition, rather it operates with the siderophore transporters in *S. lugdunensis*.

### In *S. lugdunensis* the *sst* genes are required for catecholamine-dependent iron acquisition

Catecholamine hormones enhance growth of bacteria in serum by mediating iron release from transferrin (46, 49, 51, 57). The contribution of catecholamines to the growth of *S. aureus*, via SstABCD, is only evident when endogenous SA and SB biosynthesis is eliminated (58, 59). Given that *S. lugdunensis* does not produce a known siderophore, we hypothesized that the *sst* genes may contribute significantly to iron acquisition by this bacterium. Genome analysis of *S. lugdunensis* HKU09-01 revealed the presence of two tandem *sstABCD* loci that we designated *sst*-1 and *sst*-2 that share significant similarity at the nucleotide level but that are not identical. At the protein level both Sst-1 and Sst-2 share significant identity with the SstABCD proteins in *S. aureus* (see Table S1) suggesting both loci may play a role in catecholamine transport. Analysis of *sst*-1 and *sst*-2 promoter regions identified putative Fur boxes upstream of both *sstA* genes indicating iron dependent regulation of gene expression from each *sst* locus (Figure 3A). Indeed, qPCR analysis revealed that *sst-1* is highly upregulated in iron deplete conditions as compared to iron replete conditions consistent with Fur-dependent regulation of gene expression. In contrast, expression of the *sst*-2 locus was extremely low irrespective of the iron content in the growth medium suggesting the *sst-2* locus may not function prominently in iron acquisition (Figure 3B). To begin to characterize the importance of catecholamine-dependent iron acquisition and to determine the relative contribution of the *sst-1* and *sst-2* loci to *S. lugdunensis* growth, we created a *sst* deletion mutant lacking both *sst-1* and *sst-2* (Δ*sst*1/2); for unknown reasons, we could not successfully create single *sst*-locus deletion mutants. To confirm expression of Sst proteins in *S. lugdunensis*, we performed Western blot analysis on the bacteria cultured under iron replete and deplete conditions. Using antisera generated against the *S. aureus* SstD protein we could immunodetect from wildtype *S. lugdunensis* a single protein at the expected SstD molecular weight, only under iron-deplete conditions (Fig. 3C) (46). That this anti-SstD reactive band is absent in the Δ*sst*1/2 mutant confirmed that this protein is indeed expressed from one of the Sst operons (Fig. 3C); likely SstD1 since gene expression from *sst*-2 was virtually undetectable.

**Figure 3:**
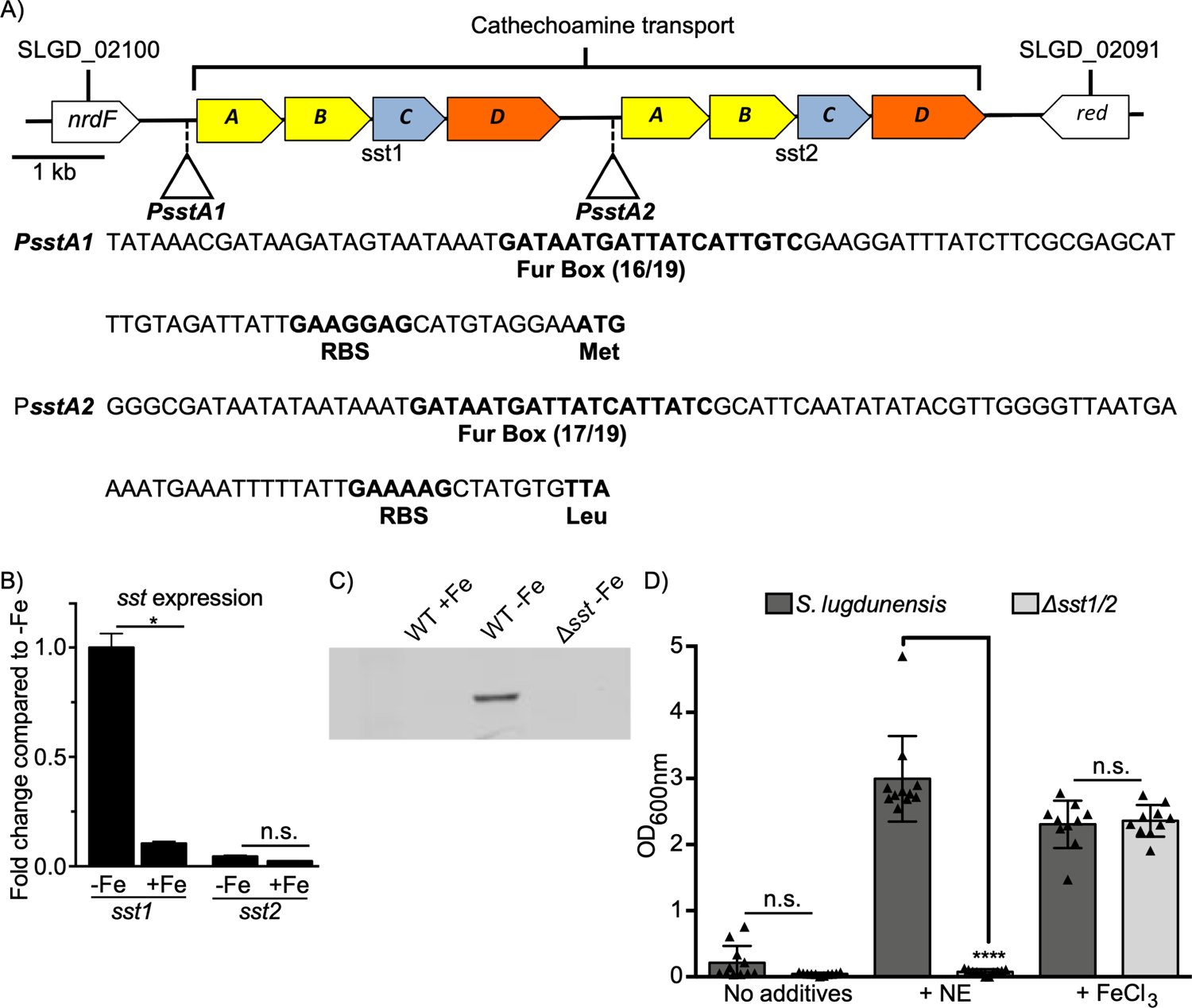
The ability of *S. lugdunensis* to utilize catecholamines for growth under iron restriction requires the *sst* genes. In (A) a physical map of putative *sst*ABCD ferric-catecholamine acquisition genes in *S. lugdunensis* is shown. The promoter sequences for each gene set are shown, with putative Fur boxes, ribosome binding sites (RBS) and start codons labeled. In *S. lugdunensis* the *sst* locus is duplicated giving rise to the operons labeled *sst*-1 and *sst*-2 respectively. In (B) the graph depicts qPCR analysis of *sst1* and *sst2* gene expression by *S. lugdunensis* grown overnight in C-TMS (-Fe) or C-TMS with 100 μM FeCl_3_ (+Fe). Data were normalized relative to *rpoB* expression, and expression was normalized relative to *sst1* in C-TMS without added iron (set as 1) as the comparator. The data derive from three independent experiments and a total of three biological replicates and statistical significance was determined by students t-test. In (C) a representative Western blot demonstrating iron-regulated expression of SstD1 in WT *S. lugdunensis* and a Δ*sst*-1/2 mutant is shown. Cultures were grown overnight in C-TMS (-Fe) or C-TMS with addition of 100 μM FeCl_3_ (+Fe). Antisera raised against *S. aureus* SstD was used to immunodetect SstD (38 kDa) from *S. lugdunensis*. In (D) the ability of *sst*-1/2 deficient *S. lugdunensis* to utilize norepinephrine (NE) as an iron source as compared to wildtype is shown. The bacteria were grown in RPMI with 1% (v/v) casamino acids and 0.05% (v/v) heat inactivated horse serum. NE was added at 50 μM and FeCl_3_ was used as a control at 20 μM. The data shown are the mean ± standard deviation of the endpoint optical density at 600nnm (OD600nm) measured after 24 hr. The data derive from three independent experiments and each symbol represents a separate biological replicate. Statistical significance was determined by ordinary one-way ANOVA with a tukey’s post-test. In B and D, n.s. indicates not significant, *p<0.05, ****p<0.0001.

To evaluate the ability of the Δ*sst*1/2 mutant to utilize catecholamines for growth under iron restriction, we compared the ability of wildtype *S. lugdunensis* and the Δ*sst*1/2 mutant to grow in iron deplete medium when norepinephrine (NE) is provided as a sole source of iron (Fig. 3D). In the absence of NE, neither wildtype nor the Δ*sst*1/2 mutant could grow unless the culture medium was supplemented with FeCl_3_, which restored growth to both wildtype and mutant bacteria. In contrast, when the culture medium was supplemented with 50 μM NE only wildtype *S. lugdunensis* was capable of growth indicating one or both *sst* loci enable NE utilization as an iron source (Fig. 3D).

We next wanted to determine the relative contributions of each *sst* operon to catecholamine utilization. Given we could not create single operon deletion mutants we chose to complement the Δ*sst*-1/2 mutant with vectors carrying the individual *sst*-1 and *sst*-2 gene sets. Provision of *sst*-1 operon *in trans* to the *Δsst*-1/2 mutant restored the ability of this strain to utilize NE and other catecholamines for growth akin to wildtype *S. lugdunensis* and as compared to the vector control (Fig. 4). Surprisingly, although we could clone the *sst-2* locus in *E. coli* and *S. aureus*, we repeatedly failed to successfully introduce this plasmid into *S. lugdunensis*. Indeed, the *S. lugdunensis* transformants that were recovered always contained plasmid carrying deletions within the cloned *sst*-2 region. Therefore, as an alternative means to determine whether the *sst*-2 locus contributes to catecholamine utilization we analyzed the growth of three *S. lugdunensis* clinical isolates whereby genome analysis identified that these strains lack the sst-1 locus while retaining the native *ss*t-2 genes; indeed, of 20 clinical isolates analyzed by whole genome sequencing three were found to lack the *sst-1* locus (see Table S3). As a control we also included a clinical isolate where both *sst-1* and *sst-2* loci are intact. This analysis revealed that the *sst-1* positive isolate grew in the presence of 20 μM norepinephrine. In contrast, all three isolates that are *sst-1* deficient failed to grow in the presence of norepinephrine despite encoding *sst-2* (Fig. S5). That these strains fail to grow, due to iron insufficiency, is evident upon addition of 20 μM FeCl_3_ which restored growth to all three isolates.

**Figure 4.**
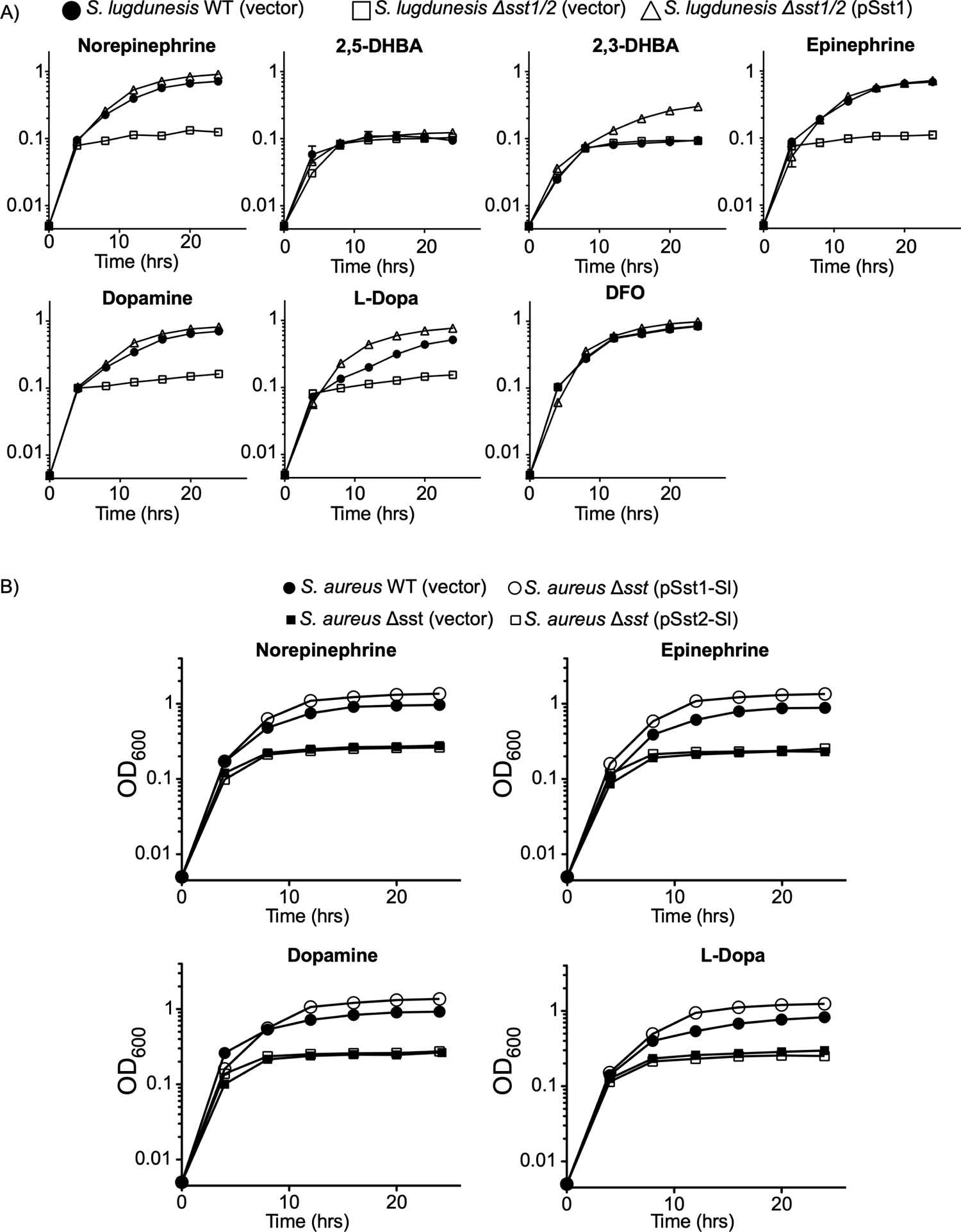
The *sst*-1 locus of *S. lugdunensis* is required for use of host catecholamine stress hormones as iron sources. In (A), growth of *S. lugdunensis* deficient for the *sst*-1/2 loci (D*sst*-1/2) was assessed in the presence of human stress hormones. Growth was compared to wildtype *S. lugdunensis* and Δ*sst*-1/2 carrying either a vector control or the p*Sst*-1 plasmid was analyzed in C-TMS with 20% (v/v) horse serum. Each catecholamine was supplemented at 50μM each catechol compound indicated and Desferrioxamine B (DFO), a hydroxamate type siderophore, was used as a control. In (B) Growth of *S. aureus sfa sbn sst* was analyzed in C-TMS with 20% serum, supplemented with 50μM of the indicated catecholamine hormones. *S. aureus* lacking *sfa* and *sbn* is labeled here as WT strain as the *sst* locus in this *S. aureus* background is intact. The strain listed as Δ*sst* carries a deletion of the *sst* locus in addition to the *sfa sbn* mutations. The bacteria were transformed with either vector control or the p*Sst*-1 or p*Sst*-2 plasmids that encode either the *sst*-1 or *sst*-2 locus from *S. lugdunensis*, respectively. Desferrioxamine B (DFO) was used as a positive control as all strains can transport the hydroxamate DFO. In (A) and (B) the data are the average of at least three independent biological replicates from three independent experiments.

In further pursuit of understanding the function of the *sst-2* locus encoded by *S. lugdunensis* we also performed heterologous complementation experiments using *S. aureus* as this staphylococcal spp. could be transformed with, and maintain, the *sst-2* expression plasmid. To assess catecholamine utilization by *S. aureus*, we utilized a *sfa sbn sst* mutant that could not synthesize endogenous SA or SB, which can obscure catecholamine uptake as previously described (46). Plasmids carrying either the *sst*-1 or *sst*-2 operon (p*sst1* and p*sst2*, respectively) from *S. lugdunensis* or the vector control were mobilized into *S. aureus sfa sbn sst* and assessed for growth under iron restricted conditions in the presence of catecholamines. As a control growth of the *sfa sbn sst* mutant was compared to *S. aureus* lacking only the *sfa* and *sbn* genes (i.e., SA and SB biosynthesis) and is referred to here as wildtype as it encodes an intact *sst* locus. In these experiments *S. aureus* lacking *sfa sbn sst* but harboring p*sst1* grew significantly better than the vector control strain in medium supplemented with norepinephrine, epinephrine, dopamine, or L-DOPA (Fig. 4B). In contrast the same *S. aureus sfa sbn sst* mutant harboring p*sst2* was impaired for growth in presence of each catecholamine and grew like the vector control. Altogether these findings indicate that the *sst*-2 locus in *S. lugdunensis* does not encode a functional catecholamine transport system however the *sst*-1 locus is both necessary and sufficient for catecholamine-iron acquisition in *S. lugdunensis*. Taken together, these observations indicate that *sst*-1 functions in iron acquisition through catecholamine utilization in *S. lugdunensis* while the function of *sst*-2 remains unknown.

### SstD proteins vary in catecholamine binding affinities

To biochemically characterize the *S. lugdunensis* SstD1 and SstD2, we investigated the substrate-binding affinities of these two proteins. To this end we overexpressed SstD1 and SstD2 from *S. lugdunensis* and SstD from *S. aureus* in *E. coli* and purified the proteins by metal-affinity chromatography. Antisera raised against *S. aureus* SstD recognized both *S. lugdunensis* SstD1 and SstD2 proteins in addition to the *S. aureus* protein (Fig. 5A). These proteins were then analyzed for substrate binding via intrinsic tryptophan fluorescence quenching. The fluorescence of all three SstD proteins was quenched in the presence of the catecholamines epinephrine, norepinephrine, dopamine, L-DOPA, and salmochelin but not in the presence of the hydroxamate DFO (Fig. 5B). The *S. aureus* SstD protein was determined to bind each catecholamine with a dissociation constant that is in agreement with a previous report (Fig. 5C) (46). The *S. lugdunensis* SstD1 protein was found to bind each of the catecholamine stress hormones analyzed, and the catecholate siderophore salmochelin, but with greater affinity than the *S. aureus* homolog (Fig. 5C). Interestingly, *S. lugdunensis* SstD2 also bound catecholamines *in vitro*, but more poorly than either of the other two SstD proteins from *S. aureus* and *S. lugdunensis*. Taken together these data reveal that the SstD proteins of *S. lugdunensis* bind to catechol compounds however based on our physiological data only the *sst1* locus contributes significantly to catecholamine dependent iron acquisition.

**Figure 5.**
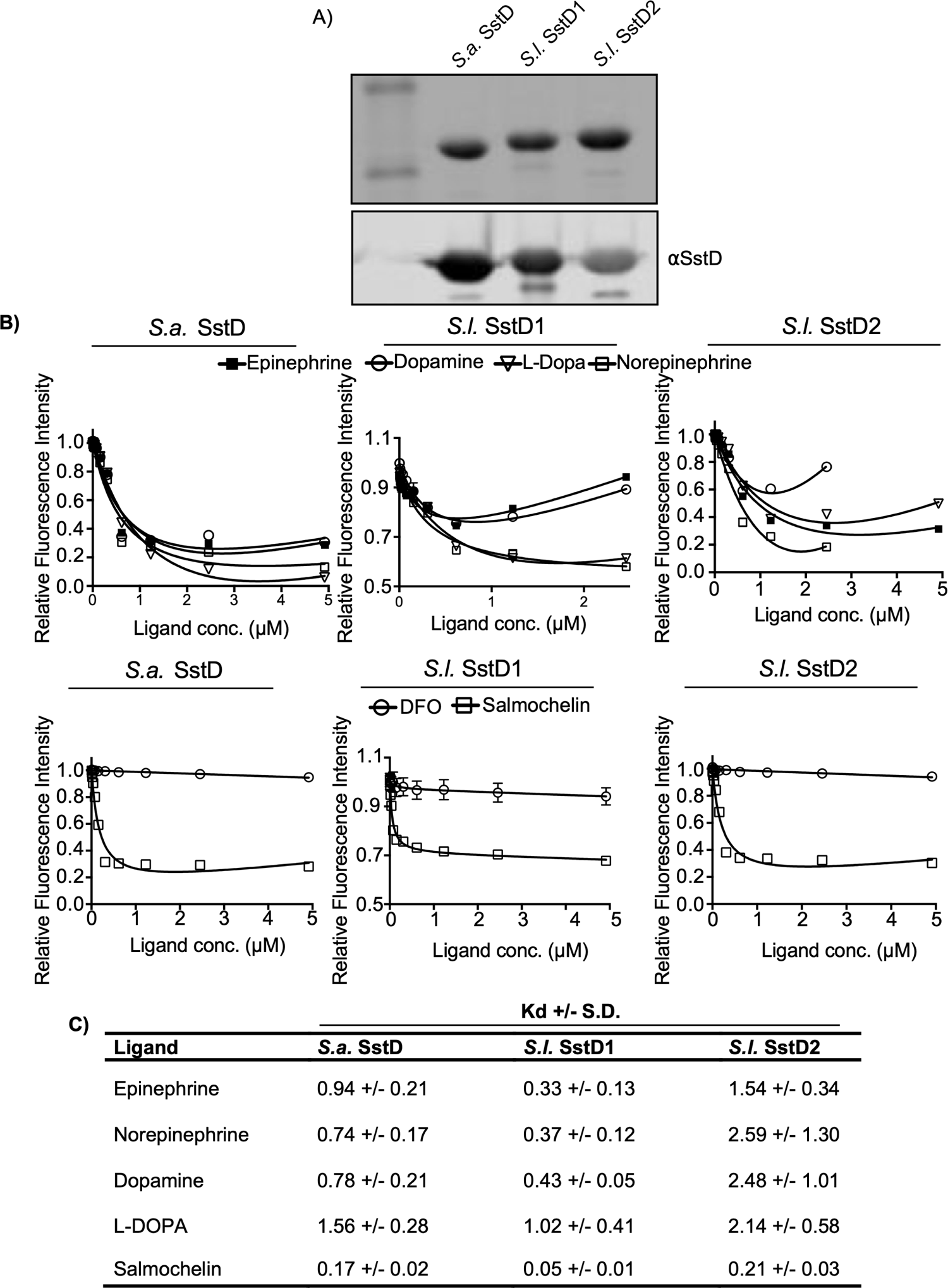
The substrate-binding properties of *S. lugdunensis* SstD1 and SstD2 proteins. In (A) a coomassie stains SDS-PAGE gel (top panel) shows the purity of the isolated recombinant SstD proteins from *S. lugdunensis* and *S. aureus*. Also shown is a representative Western blot (bottom panel) showing detection purified recombinant *S. aureus* (SA) and *S. lugdunensis* (SL) SstD homologs. Immunodetection was done with anti-SstD antiserum raised against *S. aureus* SstD protein. In (B) fluorescence quenching was used to determine binding affinity of SstD homologs for the indicated ferrated catecholamine hormones. In (C) a table listing the *K_D_* values for SstD-ferric catecholamine complexes is shown. The data are the average of at least three independent biological replicates, with error bars representing the standard error of the mean.

### Sst-dependent utilization of an iron source in casamino acids

In characterizing iron acquisition by *S. lugdunensis* experiments were performed where the bacteria were cultured in RPMI medium supplemented with 1% (w/v) casamino acids (CA). The addition of CA is frequently used to promote growth of *S. aureus* and we had previously attributed this phenomenon solely to the provision of additional amino acids. In our efforts to characterize catecholamine utilization by *S. lugdunensis* we noted that the Δ*sst*-1/2 mutant would often fail to grow in RPMI-CA. In contrast, wildtype *S. lugdunensis* grew optimally and provision of pSst-1 restored growth to the Δ*sst*-1/2 strain indicating the observed defect was indeed *sst*-1 dependent (Fig S6A). That supplementation of RPMI-CA with FeCl_3_ also corrected the Δ*sst*-1/2 growth defect demonstrated this phenotype to be iron dependent. To determine whether this observation was specific to *S. lugdunensis*, we also performed similar experiments using established mutants of *S. aureus* that lacked either *sfa sbn*, *sst* alone, or *sfa sbn* and *sst* (46). This analysis revealed that *S. aureus* also utilizes a factor that is present in CA for growth in a *sst* dependent manner and that the apparent iron acquisition defect in *S. aureus* is only evident when endogenous siderophore production is perturbed (Fig. S6B). Given that the unknown factor present in CA requires Sst catecholamine uptake systems in *S. lugdunensis* and *S. aureus* and that the growth defect is iron dependent we speculated that commercially available CA contain a catechol or related compound. To ascertain the identify of this factor, mass spectrometry was performed by two independent facilities to analyze the commercially available CA. This analysis revealed that our stock CA contained a compound or compounds related to the catecholamine methyl-DOPA, offering an explanation for the observed Sst dependent growth in RPMI when supplemented with CA.

### The ferrous iron transporter FeoAB allows *S. lugdunensis* to acquire iron at acidic pH

Analysis of the genome of *S. lugdunensis* HK09-01 revealed the presence of the transport proteins that in other bacteria have been shown to play a role in the transport of divalent metal ions including ferrous (Fe^2+^) iron (Fig. 6A). Indeed, *S. lugdunensis* carries genes encoding the FeoAB transporter (60, 61) as well as the metal ion transporter SitABC (MntABC) that, in *S. aureus*, has been shown to transport manganese (62). To characterize the contribution of *feoAB* and *sitABC* to *S. lugdunensis* growth, we created a series of deletion mutants where *feoAB* and/or *sitABC* were deleted from the bacterial genome. Growth analysis of these mutants was performed in RPMI-CA acidified to pH 5.8. In this medium *S. lugdunensis* lacking *sst*-1/2 and *fhuC* grew similarly to wildtype bacteria indicating that at pH 5.8 iron is more readily available (Fig. 6B). In contrast, a strain of *S. lugdunensis* lacking *feoAB* and *sitABC* demonstrated a modest yet statistically significant decrease in growth over a 24 hrs in the same culture medium. To ascertain whether *sitABC* or *feoAB* contributed significantly to iron acquisition at pH 5.8, these mutations were created in a Δ*fhuC* Δ*sst*-1/2 background. Growth of these mutants revealed that only when *feoAB* was inactivated, growth of *S. lugdunensis* lacking *fhuC* and *sst*-1/2 was ablated at pH5.8 (Fig. 6B). Moreover, the strains lacking *feoAB* failed to grow in an iron dependent manner as supplementation of RPMI-CA pH 5.8 with FeCl_3_ restored growth of each strain to wildtype levels (Fig. 6C). To verify the observed phenotype was indeed dependent on *feoAB*, complementation was performed and, as expected, when *feoAB* was provided in trans under the control of the native promoter growth of *S. lugdunensis* was restored to wildtype levels at pH 5.8 (Fig 6D).

**Figure 6.**
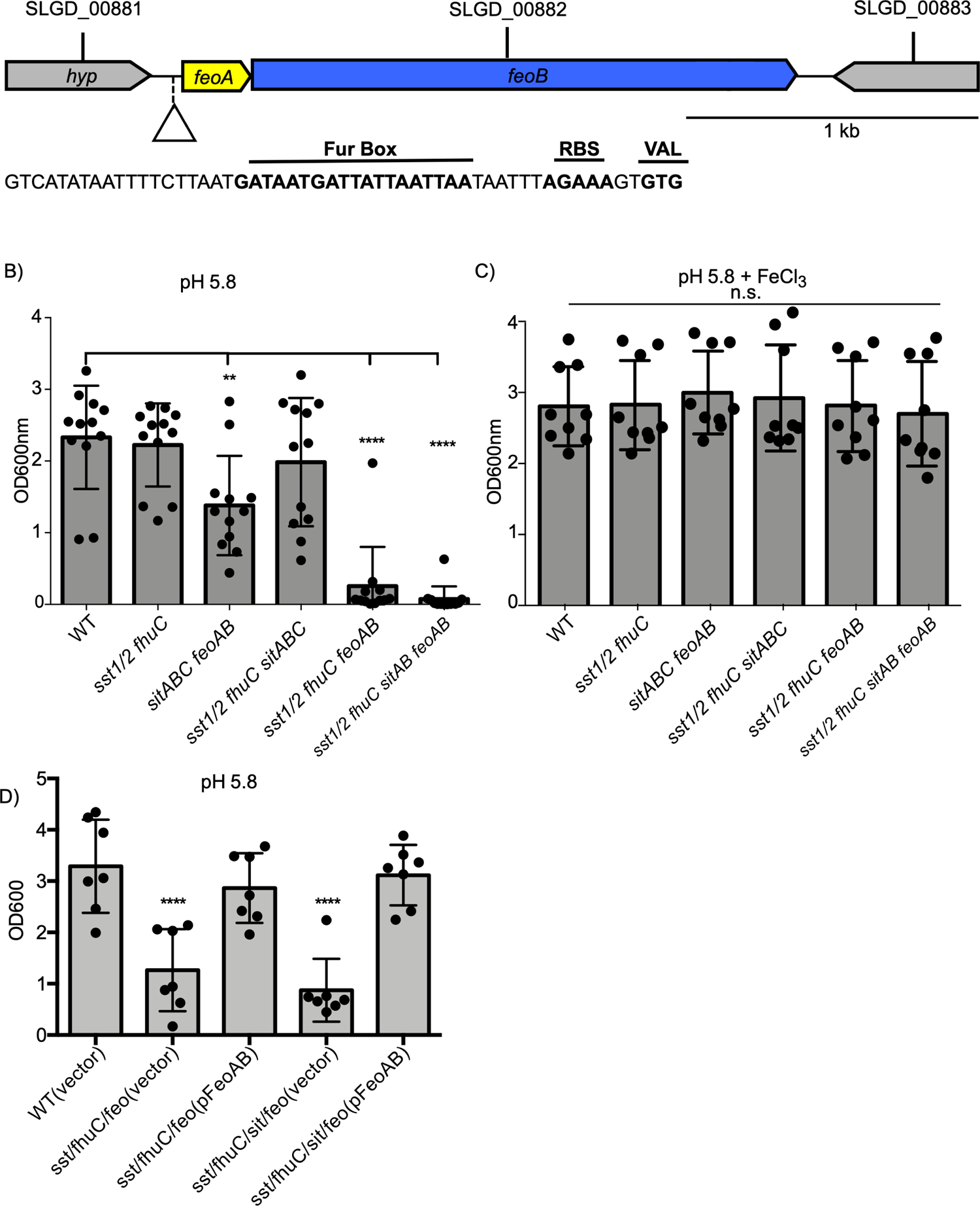
The *S. lugdunensis* genes *feoAB* that encode a putative ferrous iron transporter are required for iron-dependent growth at acidic pH. In (A) the genetic organization of the *feoAB* locus in *S. lugdunensis* is shown. The nucleic acid sequence of the upstream promoter region is also shown and the putative Fur box, ribosome binding site and alternative start codon to *feoA* is highlighted. In (B) and (C) and growth of *S. lugdunensis* mutant lacking *feoAB* in the indicated genetic backgrounds is shown. The data era the mean ± standard deviation of the measured OD600nm after 24 h. In (B) the bacteria were grown in RPMI pH5.8 whereas in (C) the bacteria were grown in the same media but supplemented with 20 μM FeCl_3_. In (D) similar growth analysis was performed except the indicated *S. lugdunensis* strains were transformed with a vector control or the p*FeoAB* plasmid. In (B, C and D) each symbol represents a biological replicate, and the data derive from three independent experiments. Statistical significance was determined by a one-way ANOVA with a Dunn’s post-test where each data set was compared to WT. n.s. indicates not significant, *p<0.01, ****p<0.0001.

### Comprehensive iron acquisition allows *S. lugdunensis* to proliferate in murine kidneys

Iron is scarcely available within the mammalian host. The preceding experiments revealed the importance of several iron acquisition systems in *S. lugdunensis* and the conditions with which they function to acquire iron *in vitro*. However, we next sought to establish their importance during infection. Previous work from our laboratories has established that *S. lugdunensis* utilizes the iron regulated surface determinant pathway (Isd) as well as the LhaSTA transporter, encoded from within the *isd* locus, to acquire iron from heme and hemoglobin (36, 53). Given hemoglobin/heme are relevant sources of iron *in vivo* we examined phenotypes for the isd mutant and, as shown in Fig. 1, found that bacterial burdens in kidneys were no different than for that of mice infected with wild type bacteria. Thus, other iron acquisitions mechanisms must be at play. Combining mutations in *sst and fhuC* into the *isd* mutant (the mutant had a confirmed inability to grow on hemin as a source of iron, Fig. S2), yielded a strain that was attenuated in the first several days of infection but eventually the bacterial burden reached levels similar to those seen in wild type-infected mice (Fig. 7A). These data indicated that while *fhuC*, *sst,* and *isd* were involved in the early stages of infection in the kidneys, other iron acquisition systems must also function to allow the bacteria to eventually grow. Importantly however, these data indicate that *S. lugdunensis* must initially utilize catechol and/or hydroxamate type iron chelates that must exist within the host. To elucidate whether the *feo* and/or *sit* genes contribute to growth in vivo we next incorporated mutations in these loci in the *fhuC sst isd* background and observed that this mutant was now attenuated (>2 log relative to wild type) for proliferation in the kidneys through day 7 of the infection. In strains lacking mutations in either isd or *feo*/*sit*, there was no significant attenuation, indicating combined disruption of all of *isd*/*fhuC*/*sst*/*feo*/*sit* was required to establish long-lasting perturbation of infection in mice. Taken together these data indicate that the *isd* genes in addition to the non-heme iron acquisition systems must operate in *S. lugdunensis* for growth within murine kidneys.

**Figure 7.**
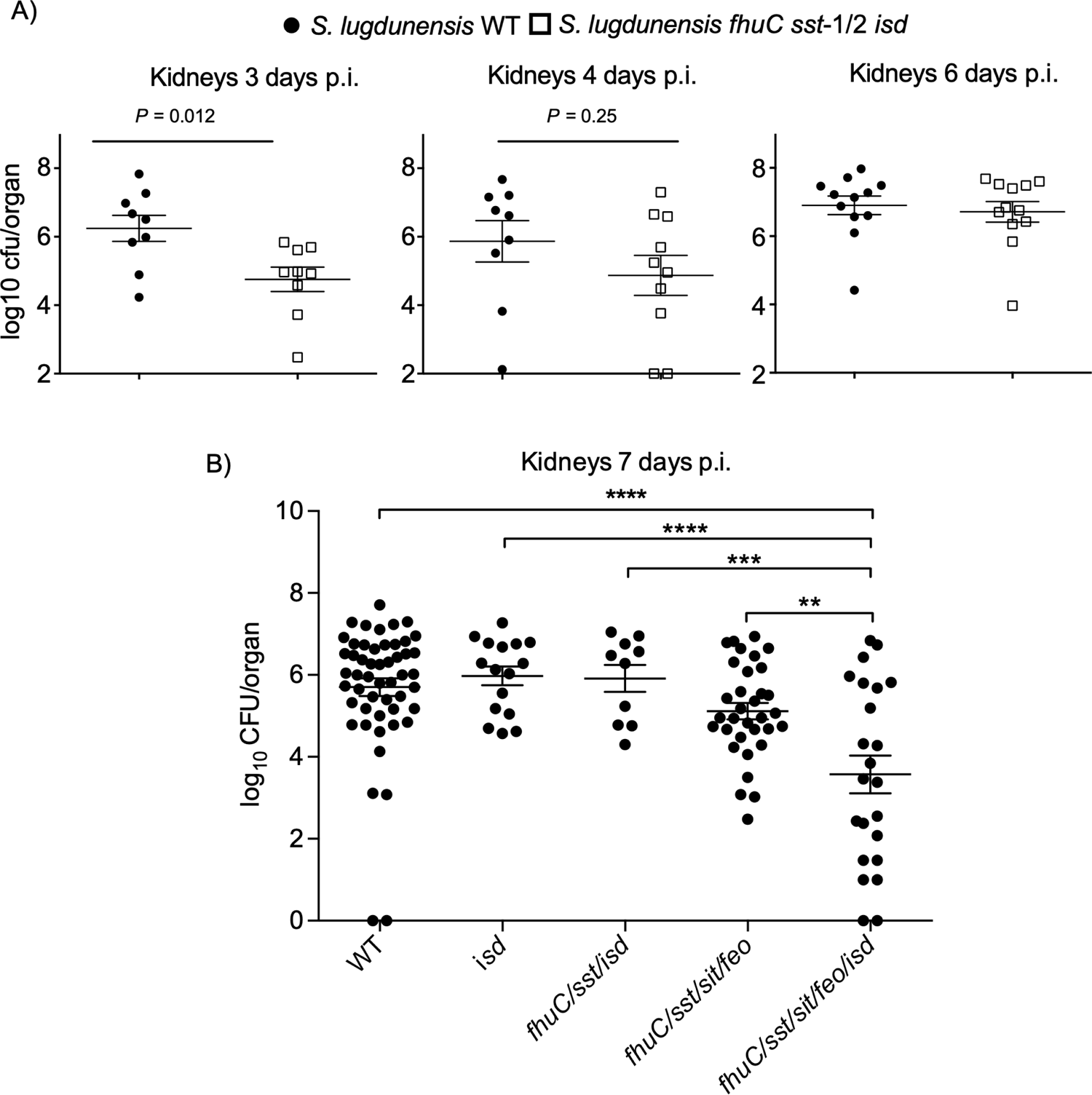
Disruption of all of Isd, FhuC, Sst, Sit and Feo in *S. lugdunensis* is required to attenuate bacterial growth in murine kidneys. Female BALB/c mice were infected systemically with 2-3×10^7^ CFU *S. lugdunensis* wildtype or an isogenic *isd fhuC sst* mutant. In (A) the bacterial burden in the kidneys of infected mice is shown. Mice were sacrificed at day 3, 4 and 6 and each symbol represents the measured burden expressed as log10 CFU from both kidneys of a single mouse. The mean is depicted the horizontal bar and the error bars represent the standard error of the mean for each group. Statistical analyses were performed using an unpaired Student’s *t*-test. The limit of detection is the y-axis value at the origin. In (B) the burden of *S. lugdunensis* strains lacking combinations of iron acquisition systems in the murine kidney at day 8 is shown. The data are presented as the mean log10 CFU/organ where the horizontal bar is the mean and the bars are the standard deviation. Each symbol represents a single animal. Statistical significance was measured using an ordinary one-way ANOVA using a tukey’s multiple comparison test. The horizontal bars indicate comparisons where statistically significant differences are observed and **p<0.01, ***p<0.001, ****p<0.0001.

## DISCUSSION

The capacity to acquire iron underpins the ability of bacteria to cause infection and *S. lugdunensis* thrives in diverse niches within the host where it can cause a spectrum of diseases. The ability of *S. lugdunensis* to cause disease necessitates this bacterium must deploy iron acquisition systems, however, unlike other staphylococci such as *S. aureus* it does not produce a siderophore. Nevertheless, *S. lugdunensis* encodes within its genome both heme and non-heme dependent iron acquisition systems that allow for proliferation when confronted with an intact mammalian immune system. This is evidenced by the observation that the burden of *S. lugdunensis* within the murine kidney increases over time (see Fig. 1C) and that mutagenesis of iron acquisition genes can antagonize *S. lugdunensis* infection (see Fig. 7A and B). Our infection data demonstrate that while inactivation of the Isd system alone in *S. lugdunensis* attenuates growth on hemin *in vitro*, Isd mutagenesis does not attenuate growth in the murine kidney. Ostensibly, this is because the IsdB protein of *S. lugdunensis* binds murine hemoglobin with reduced affinity as compared to human hemoglobin (35) and *S. lugdunensis* demonstrates weak hemolytic activity toward murine erythrocytes (37). Nonetheless, the underlying importance of the Isd pathway is evident when additional iron acquisition genes such as *fhuC, sst*-1/2, and *feoAB* are inactivated. Conversely, that an *isd* mutant alone also does not present with any defects *in vivo* as compared to wildtype, indicates that these other non-heme iron acquisition systems must also operate *in vivo*. Collectively the Fe acquisition pathways that function in *S. lugdunensis* must work in a concerted manner to provide sufficient iron to support *S. lugdunensis* growth in the murine kidney. Moreover, due to their overlapping function in metal acquisition, multiple mutations are required for *in vivo* phenotypes to be evident. Ostensibly, this is because many different sources of iron exist, and functional redundancy ensures that the bacteria can access this critical metal within the host.

Based upon *in vitro* analyses, the importance of *S. lugdunensis* Fe acquisition using only specific iron sources has enabled us to demonstrate that this bacterium can transport a variety of siderophores, mammalian stress hormones and even ferrous iron. That *S. lugdunensis* cannot synthesize a siderophore distinguishes this *Staphylococcus* spp. from *S. aureus* and other CoNS (44). Moreover, the inability of *S. lugdunensis* to synthesize siderophores explains why this bacterium is incapacitated for growth even in very low concentrations of horse serum (a source of transferrin) *in vitro* (see Fig. 1A and (44)). Despite not synthesizing siderophore, *S. lugdunensis* can usurp xenosiderophores (i.e., siderophores produced by other bacteria). Indeed, *S. lugdunensis* can utilize hydroxamate-type siderophores such as aerobactin produced by *E. coli* and polycarboxylate siderophores such as SA and SB, synthesized by *S. aureus* (see Fig. S3 and (44)). *S. lugdunensis* can also utilize catecholamine stress hormones that can interact with holotransferrin and liberate the bound iron rendering it available to the bacteria (52). Catecholamines have been shown to promote growth of many other bacteria (52, 57) in addition to *S. aureus* under iron limiting conditions. However, in the case of latter, the importance of catecholamine-dependent iron acquisition is only evident in the absence of endogenous staphyloferrin production (46). It is interesting that the *sfaA* and *sfaD* genes involved in SA biosynthesis and export are deleted in *S. lugdunensis* (44). Presumably, the inability to synthesize SA has unburdened *S. lugdunensis* with the need to consume metabolites and expend energy for siderophore biosynthesis but why this would be of benefit to *S. lugdunensis* is unclear as the bacteria would be reliant on exogenous iron chelates.

*S. lugdunensis* strain HKU09-01 differs from *S. aureus* in that it encodes two tandem *sst* loci (*sst*-1 and *sst*-2) both of which were predicted to function in catecholamine use; however, our experiments clearly demonstrated that the ability of *S. lugdunensis* to derive iron from catecholamines can be attributed solely to the *sst*-1 locus. *S. lugdunensis* is unique among staphylococci in that it can carry duplicated *sst* gene sets, and this distribution of putative *sst* genes is seen in many clinical isolates (31, 32). It is interesting that three clinical isolates were identified that lack the *sst-1* locus however as our data reveal these strains fail to utilize NE as an iron source and it is unclear why such gene loss would occur in a background where siderophore is already not produced. Despite sst-1 clearly playing a role in catecholamine utilization, we fail to detect expression or show a function for *sst*-2 in catecholamine utilization or otherwise. That the SstD2 protein of *S. lugdunensis*, as compared to the other SstD homologs, displays reduced affinity for catecholamine substrates could suggest that the second *sst* locus in *S. lugdunensis* could transport a different nutrient altogether. At present, it is unknown what purpose the *sst*-2 locus serves for *S. lugdunensis*. In contrast, our data clearly show that *sst*-1 allows this bacterium to utilize catecholamines and perhaps in the absence of endogenous siderophore production, *S. lugdunensis* is often reliant on catechols for growth during infection as compared to *S. aureus*. Indeed, *S. lugdunensis* expresses SstD-1 which has a greater affinity for catechols than the *S. aureus* SstD ortholog. It is noteworthy that in *S. lugdunensis* the IsdB protein, a component of the Isd pathway, was also found to bind hemoglobin with greater affinity than its counterpart in *S. aureus* (35, 63). Conceivably, as *S. lugdunensis* lacks the ability to synthesize SA or SB, unlike *S. aureus*, this bacterium has evolved optimized SA and SB independent iron transport systems to compensate.

That *S. lugdunensis* cannot make siderophore might also render this bacterium more reliant on other siderophore-independent metal transporters such as FeoAB. Feo transporters function to transport ferrous iron into bacteria and these systems have been better characterized in Gram-negative organisms (61). In contrast, several Gram-positive bacteria encode Fur-regulated Feo transport systems, however, their role in iron acquisition has remained largely uncharacterized (23). Here we show for the first time for a *Staphylococcus* spp. that FeoAB are required for growth at acidic pH in an iron-dependent manner. Under these conditions (i.e., pH 5.8) iron should exist in the Fe^2+^ state and *in vivo* Fe^2+^ might exist in hypoxic/anoxic or acidic environments such as abscesses or phagolysosomes within immune cells. In contrast to FeoAB, a role for the *sitABC/mntABC* locus in iron acquisition in *S. lugdunensis* could not be found *in vitro* indicating FeoAB is primarily utilized by this organism for Fe^2+^ utilization.

During our investigation, we found that *S. lugdunensis* displayed improved growth in the presence of casamino acids used to supplement RPMI. Naively, we initially attributed the improved growth of *S. lugdunensis* to the provision of additional amino acids, however, we found that improved growth of *S. lugdunensis* in RPMI with CAS was dependent on the *sst*-1 locus. Moreover, compromised growth of a *sst* mutant could be rescued by the addition of iron not amino acids. These observations indicated that CAS provided the bacteria with an additional iron source and mass spectrophotometry confirmed that catecholamines, such as methyl-dopa, can be present in commercial CAS preparations. As CAS is often used to supplement bacterial media such as tris-minimal succinate or RPMI (46, 64, 65), caution should be taken as impurities present in CAS could have unintended effects on experimental outcome.

*S. lugdunensis* lacks the vast arsenal of virulence factors present in *S. aureus*, which may explain why the bacterial burden in the liver of infected animals decreases with time. In contrast, the burden of *S. lugdunensis* does increase in the kidney over time (see Fig. 1C). Previously, it has been shown that a heme auxotroph of *S. aureus* is better able to replicate within the murine kidney, as compared to other visceral organs, suggesting the kidney might be a niche where heme is more readily availability (54). Therefore, *S. lugdunensis* might be poised to grow in this organ as it expresses both Isd and an energy-coupling factor (ECF) type ABC transporter specific for heme (37, 44, 53). Both systems are Fur-regulated and therefore expressed under iron limiting conditions (i.e., *in vivo*), however other low affinity heme transport systems might also exist. Interestingly, the Isd proteins and ECF heme transporter in *S. lugdunensis* enable growth on heme of murine and human origin, however, the hemolytic factors secreted by *S. lugdunensis* have a propensity to lyse only human erythrocytes (37). Indeed, studies have reported that in a systemic murine model of infection *S. lugdunensis* fails to cause significant morbidity (66, 67), an observation corroborated here, which could be in part be attributable to the presence of human specific hemolysins. Nevertheless, animal models are an important tool that can be employed to study *S. lugdunensis* growth and metal acquisition in the mammalian host and that *S. lugdunensis* failed to cause significant morbidity facilitated our study. Indeed, through *in vivo* infection experiments we have demonstrated that several genetic loci within *S. lugdunensis* allow the bacteria to procure iron from an assortment of non-heme and heme related sources *in vitro*, however, *in vivo* it is the concerted action of these systems that allows *S. lugdunensis* to grow. Therefore, the development of interventions that target bacterial iron acquisition systems should consider the overlapping function of distinct metal acquisition strategies deployed by bacterial pathogens.

### Experimental procedures

#### Bacterial strains and media

Bacterial strains and vectors employed in this study are summarized in Table 1. *Escherichia coli* strains were grown in Luria-Bertani broth (LB, BD Diagnostics) or on LB agar. For routine culture and genetic manipulation, *S. lugdunensis* and *S. aureus* strains were cultured in tryptic soy broth (TSB) or on TSB solidified with 1.5% (w/v) agar (TSA). For growth experiments *S. lugdunensis* and *S. aureus* were cultured in RPMI 1640 (Life Technologies) that was in some instances supplemented with 1% (w/v) casamino acids (BD Diagnostics) (RPMI-C). Growth was also performed in tris-minimal succinate (TMS) broth or 1.5% (w/v) agar (68). TMS broth was treated with 5% (w/v) Chelex-100 resin (Bio-Rad) at 4°C for 24 hours (C-TMS) to chelate trace metals. As appropriate, the above media were supplemented with heat-inactivated horse serum (Sigma Aldrich) which served as a source of transferrin to restrict iron availability. Bacteria were cultured at 37°C with shaking at 220 rpm unless otherwise indicated. For *E.* coli antibiotic selection was as follows: 100 µg mL^-1^ ampicillin or 50 µg mL^-1^ kanamycin. For *S. lugdunensis* and *S. aureus* antibiotic selection was as follows: 10 to 12 µg mL^-1^ chloramphenicol, 4 µg mL^-1^ tetracycline, and 50 µg mL^-1^ kanamycin.

#### Real-time PCR

Quantitative real-time PCR was performed as previously described (69). Briefly, *S. lugdunensis* HKU09-01 RNA was prepared from triplicate 3 mL cultures grown in C-TMS or C-TMS with 100 µM FeCl_3_. Cultures were harvested to an OD_600_ of 3.0 and RNA was extracted using the Aurum Total RNA Mini Kit (BioRad). 500 ng extracted RNA was reverse-transcribed and PCR-amplified using iScript^TM^ One-Step RT-PCR Kit with SYBR Green (Bio-Rad) and primers outlined in Table 1. Data were normalized relative to expression of the *rpoB* housekeeping gene.

#### Gene deletion and complementation of *S. lugdunensis*

For in-frame deletion of *S. lugdunensis* genes allelic replacement using the pKOR1 or pIMAY vector was performed as described previously (70, 71). Briefly, 500 – 1,000-bp DNA fragments flanking regions of interest were amplified using the primers found in Table 1. Upstream and downstream flanking amplicons were cloned into pKOR1 or pIMAY. Knockout vectors were passed through *S. aureus* RN4220 or *E. coli* SL01B before introduction into *S. lugdunensis* by electroporation (66, 71). Plasmids were integrated into the genome at 42°C for pKOR1 or at 37°C for pIMAY in the presence of chloramphenicol prior to counter-selection at 30°C in the presence of anhydrotetracycline (pKOR1: 200 ng mL^-1^, pIMAY: 1 µg mL^-1^). Chloramphenicol-sensitive colonies were chosen for screening by PCR across the deleted region in the chromosome, which was further confirmed by sequencing (70, 71). The same process was used over to generate multiple deletions in one strain.

For complementation, the *S. lugdunensis fhuC, sstA1B1C1D1* and *sstA2B2C2D2* and feoAB genes were PCR amplified using primers described in Table 1 and each amplicon encompassed the native upstream promoter. The *fhuC, sstA1B1C1D1* and *sstA2B2C2D2* amplicons were cloned into the plasmid pRMC2 to create of p*fhuC*, p*sst1* and p*sst2*. The *feoAB* amplicon was cloned into the plasmid pALC2073. Each amplicon was cloned as a KpnI/SacI fragment and plasmids were confirmed by DNA. All cloning was performed in *E. coli* DH5α and the resulting plasmids were passed through *S. aureus* RN4220 or *E. coli* SLO1 prior to electroporation into the appropriate *S. lugdunensis* mutant.

### Genome sequencing

*S. lugdunensis* clinical isolate genome sequencing data are deposited under accession PRJNA796272.

### Siderophore preparation and plate bioassays

*S. aureus* concentrated culture supernatants were prepared from Δ*sbn,* Δ*sfa,* and Δ*sbn*Δ*sfa* mutants, respectively, as described previously (39). Strains were grown in C-TMS with aeration for 36 hours prior to removal of cells. Supernatants were lyophilized and insoluble matter was removed by methanol extraction (one-fifth original culture volume). Methanol was removed by rotary evaporation, and dried material was resuspended in water to one-tenth culture volume to provide culture extracts. Staphyloferrin B was prepared *in-vitro* enzymatically, as described previously (40, 43, 72). Enzymes were removed from the reaction mixture using an Amicon Ultra-0.5 10k filter column (Millipore) and the Staphyloferrin B reaction mixture was normalized to Deferoxamine (DFO, London Health Sciences Center) equivalents as determined using the chrome azurol S (CAS) siderophore detection assay (73). Staphyloferrin A was commercially prepared by Indus BioSciences (India). Ferric–enterobactin, -salmochelin S4, -aerobactin and coprogen were purchased from EMC Microcollections. Ferrichrome was purchased from Sigma, whereas citrate was purchased from Fisher Scientific.

The ability of culture supernatants and purified siderophores to support *S. lugdunensis* iron-restricted growth was assessed with agar plates using plate-based disk diffusion bioassays (44, 45). Briefly, 1 x 10^4^ *S. lugdunensis* cells were incorporated into TMS-agar containing 5 µM ethylenediamine-di(*o*-hydroxyphenylacetic acid) (EDDHA, LGC Standards GmbH). Siderophores/supernatants applied to sterile paper disks were placed onto the agar, and growth around disks was measured after 24 hours at 37°C.

### SstD Western blot analysis

Antisera against *S. aureus* SstD, used in this study, was previously prepared (46) and was used for analysis of iron-regulated SstD expression in *S. lugdunensis*. *S. lugdunensis* bacteria were grown in C-TMS with or without 100µM FeCl_3_ for 24 hours, normalized and lysed in the presence of lysostaphin (Sigma). Whole cell lysates were normalized to 8 µg total protein and resolved by SDS-polyacrylamide gel electrophoresis. Western blotting was performed as previously described (44). In brief, the membrane was blocked in phosphate buffered saline (PBS) with 10% (w/v) skim milk, 0.05% (v/v) Tween 20 and 20% (v/v) horse serum. Antiserum was applied at a 1:5,000 dilution in PBS with 0.05% Tween 20 and 5% horse serum, prior to addition of anti-rabbit IgG conjugated to IRDye-800 (1:20,000 dilution; Li-Cor Biosciences). Fluorescence imaging was performed using a Li-Cor Odyssey infrared imager (Li-Cor Biosciences).

### Growth in serum

Growth of *S. lugdunensis* and *S. aureus* strains was assessed in C-TMS with serum. Single, isolated colonies were resuspended in 2 mL C-TMS and grown for over 4 hours until OD_600_ was above 1. Each culture was normalized to an OD_600_ of 1 and subcultured 1:200 in C-TMS:horse serum. Wildtype *S. lugdunensis* as well as *S. aureus* strains bearing Δ*sbn* and Δ*sfa* mutations, are impaired for growth in this media compared to siderophore-producing strains (39, 44). Human stress hormones were added to the media for a final concentration of 50 µM to assess for catecholamine-iron acquisition for growth enhancement. Dopamine hydrochloride, L-3,4-dihydroxyphenylalanine (L-DOPA), DL-norepinephrine hydrochloride, (-)-epinephrine, 2,3-dihydroxybenzoic acid (DHBA) and 2,5-DHBA were purchased from Sigma. Chloramphenicol was also included for strains harboring pRMC2 or derivatives. Cultures were grown in a Bioscreen C plate reader (Growth Curves USA) at 37°C with constant shaking at medium amplitude. OD_600_ was assessed at 15-minute intervals however, for graphical clarity, 4 hour intervals are shown.

### Protein overexpression and purification

Recombinant *S. aureus* SstD was purified as previously described (46). Regions of the genes encoding the soluble portions of *S. lugdunensis* SstD1 and SstD2 (excluding lipobox motifs) were amplified and cloned into pET28(a)+ (Novagen) using primers listed in Table 1. *E. coli* BL21 bearing pET28::*sstD1* or pET28::*sstD2* were grown to mid-log phase at 37°C in LB with kanamycin, prior to induction with 0.4 mM isopropyl-β-D-1-thiogalactopyranoside (IPTG). After addition of IPTG, cultures were grown at 25°C overnight. Cells were collected by centrifugation, resuspended in 20 mM Tris, pH 8.0, 500 mM NaCl, 10 mM imidazole (binding buffer), and ruptured in a cell disruptor (Constant Systems Ltd). Insoluble matter and debris were removed by centrifugation at 3,000 x g for 15 minutes, followed by 150,000 x g for 60 minutes, sonicating samples in between. Soluble material was filtered and applied to a nickel-loaded 1 mL HisTrap column (GE Healthcare) equilibrated with binding buffer. His_6_-tagged proteins were eluted in 1 mL fractions from the column over a 0-80% gradient of 20 mM Tris, pH 8.0, 500 mM NaCl, 500 mM imidazole (elution buffer). Fractions bearing pure SstD1 and SstD2 (analyzed via SDS-PAGE) were pooled and dialyzed into 10 mM Tris, pH 8.0, 100 mM NaCl (working buffer) at 4°C. Protein concentrations (Bio-Rad protein assay) were normalized to equality and aliquots were frozen at −80°C.

### Protein-ligand binding

Intrinsic tryptophan fluorescence quenching was used to assess protein-ligand binding affinity for *S. lugdunensis* SstD1, SstD2 and *S. aureus* SstD as previously described (46). Proteins were adjusted to 0.5 μM in 3 mL working buffer and ligands were added at 2-fold concentration increments. Dopamine, L-DOPA, epinephrine, norepinephrine, DFO and salmochelin S4 were used as ligands. Ligands were incubated in 3:1 (catecholamine hormones) or 1:1 (siderophores) molar ratio to FeCl_3_ for 5 minutes at room temperature prior to use. Bovine serum albumin (Sigma) was used as a protein negative control. Fluorescence was measured at room temperature in a Fluorolog instrument (Horiba Group), with excitation at 280 nm and emission detection at 345 nm. An excitation slit width of 5 nm and an emission slit width of 5 nm were used. Changes in fluorescence due to ligand additions and sample volume increase were corrected for (74). Fluorescence intensity data analysis and *K_D_* determination were performed as previously described (46).

### Analysis of recombinant protein expression

Recombinant proteins were analyzed for purity and immunogenicity towards αSstD (*S. aureus*) antisera. *S. lugdunensis* SstD1, SstD2 and *S. aureus* SstD purified protein volumes were normalized to contain 3 μg total protein and resolved by SDS-PAGE. Western blotting was performed as described above with the following modifications. After blocking, αSstD antisera were applied at a 1:20,000 dilution, and αHis antibody was applied 1:10,000. Anti-rabbit IgG conjugated to IRDye-800 (1:20,000 dilution) was secondary to αSstD antisera, whereas anti-mouse Alexa Fluor 680 (Life Technologies) was secondary to αHis (1:20,000 dilution). Antibodies/antisera were applied in PBS with 0.05% Tween 20 and 5% horse serum.

### Analysis of *S. lugdunensis* growth under iron restriction

*S. lugdunensis* with or without plasmids were grown O/N at 37 °C on TSA plates in the presence of selection as appropriate. Isolated colonies were resuspended in 2 mL of growth medium (i.e., C-TMS or RPMI) in 14 mL polypropylene snap cap tubes and grown overnight at 37 °C with shaking at 225 rpm. Each culture was pelleted and washed twice in sterile 0.9% (w/v) saline and normalized to an OD_600_ of 0.5. Next day 2 mL cultures were set up in 14 mL polypropylene snap cap tubes containing RPMI, RPMI-C or RPMI at pH5.8 (acidified with HCl) with or without horse serum as necessary. Cultures were inoculated at a starting OD_600_ of 0.005 and were grown for 18-24 h at 37C with shaking at 225 rpm. Endpoint OD600 was read to evaluate the ability of *S. lugdunensis* to grow. In some instances, the additive DFO (100 μM), norepinephrine (NE, 50 μM), or hemin (50 nM) or FeCl_3_ (20 μM) was added to some cultures.

Growth curves were monitored using a BioScreen C plate reader with constant shaking at medium amplitude, at 37°C. OD_600_ was measured at 15-minute intervals and growth at 4-hour intervals are shown. Here growth of *S. lugdunensis* and *S. aureus* strains was assessed in C-TMS with serum. Isolated colonies were resuspended in 2 mL C-TMS and grown for over 4 hours until OD_600_ was above 1. Each culture was normalized to an OD_600_ of 1 and subcultured 1:200 in C-TMS:horse serum. Cultures were pipetted into BioScreen C honeycomb plates in 200 μL culture volumes. Wildtype *S. lugdunensis* as well as *S. aureus* strains bearing Δ*sbn* and Δ*sfa* mutations, are impaired for growth in this media compared to siderophore-producing strains (39, 44). Human stress hormones were added to the media for a final concentration of 50 µM to assess for catecholamine-iron acquisition for growth enhancement. Dopamine hydrochloride, L-3,4-dihydroxyphenylalanine (L-DOPA), DL-norepinephrine hydrochloride, (-)-epinephrine, 2,3-dihydroxybenzoic acid (DHBA) and 2,5-DHBA were purchased from Sigma.

### Murine model of systemic *S. lugdunensis* infection

All protocols for murine infection were reviewed and approved by the University of Western Ontario’s Animal Use Subcommittee, a subcommittee of the University Council on Animal Care. Six-week-old, female, BALB/c mice were obtained from Charles River Laboratories and housed in microisolator cages. *S. lugdunensis* strains were grown to mid-exponential phase (OD_600_ 2 - 2.5) in 25 mL TSB, washed twice with PBS, and resuspended in PBS to an OD_600_ of 0.50. Next, 100 µL of bacterial suspension, equivalent to ∼2-3 x 10^7^ CFU, was injected into each mouse via tail-vein. Mice were weighed at time of challenge and every 24 hours after, where infection was allowed to proceed for 3 days before mice were euthanized via cervical dislocation. Organs were aseptically harvested into 3 mL PBS with 0.1% (v/v) Triton X-100, homogenized, diluted, and plated onto TSA to enumerate bacterial burden. Weight data are presented as the difference in percentage from mouse weight at time of challenge. Recovered bacterial load from organs is presented as log_10_ CFU per organ.

### Human hemoglobin purification

Human hemoglobin was purified as described elsewhere (75).

### Analysis of *S. lugdunensis* growth with human hemoglobin

*S. lugdunensis* N920143 WT, Δ*isdL*, Δ*fhuC* and Δ*isdL*Δ*fhuC* was grown O/N in TSB at 37°C with shaking at 160 rpm. Cells were pelleted, washed with RPMI supplemented with 1% casamino acids and 10 µM EDDHA, and adjusted to OD_600_ = 1. 2.5 µl of these cultures were used to inoculate 500 µl of RPMI + 1% casamino acids + 10 µM EDDHA per well (starting OD_600_ of 0.005) in a 48 well microtiter plate (Nunc, Thermo Scientific). As iron sources, 2.5 µg/ml human hemoglobin (hHb, own preparation) or 20 µM FeSO_4_ (Sigma-Aldrich) were added. Growth was measured using an Epoch2 reader (BioTek) (37°C, orbital shaking) every 15 min for 48 hrs.

### Plasmid constructions for BACTH

*S. lugdunensis* N920143 WT chromosomal DNA was used to amplify *isdF*, *isdL*, and *fhuC*, and the fragments were cloned into the vectors pKT25 and pUT18C (Euromedex), respectively, by restriction digestion. Used primers can be found in Table 1. After transformation into *E. coli* XL-1 blue, colonies were confirmed by sequencing, Bacterial adenylate cyclase two-hybrid system (BACTH) assay To investigate interaction between the permease IsdF and the ATPases IsdL and FhuC, the commercially available BACTH kit was used (Euromedex). In brief, *E. coli* BTH101 was co-transformed with pKT25:*isdF* and pUT18C:*isdL* or pUT18C:*fhuC*, respectively. In case of protein-protein interaction, the catalytic domains T25 and T18 of the *Bordetella pertussis* adenylate cyclase are able to heterodimerize and to produce cyclic AMP (cAMP) allowing the expression of *lacZ*. This leads to blue colony formation on LB agar indicator plates containing 40 µg/mL X-Gal (Sigma-Aldrich/Merck), 0.5 mM isopropyl β-D-1-thiogalactopyranoside (IPTG) (Thermo Scientific), 100 µg/mL ampicillin and 50 µg/mL kanamycin after incubation for 2 days at 30°C. As positive control, pKT25:*zip* and pUT18C:*zip* were used encoding a leucine zipper; as negative control, empty vectors were co-transformed into BTH101.

## ACKNOWLEDGEMENTS

This work was supported by an operating grant to DEH from the Natural Sciences and Engineering Research Council (NSERC). SH acknowledges funding by the Deutsche Forschungsgemeinschaft (DFG) from an individual project grant (HE8381/3-1). SH was supported by infrastructural funding from the Deutsche Forschungsgemeinschaft (DFG), Cluster of Excellence EXC 2124 Controlling Microbes to Fight Infections. JRB was supported by a Queen Elizabeth II Graduate Scholarship in Science and Technology.

---

**Figure S1. Analysis of *S. lugdunensis* burden in visceral organs during systemic murine infection.** Mice were infected with wild-type *S. lugdunensis* and the bacterial burden in the indicated organs was determined 3 days (72 h) post-infection where each symbol represents an individual animal. The data presented are the mean ± standard deviation for the determined log 10 CFU/organ. Statistical significance was determined using an ordinary one-way ANOVA with a tukeys multiple comparison test. *p<0.05, **p<0.01.

**Figure S2. *S. lugdunensis* lacking the *isd* locus cannot utilize hemin as an iron source.** Wild-type *S. lugdunensis* HKU09-01 and mutants lacking *isd* were cultured in RPMI supplemented with 0.1% (v/v) horse serum. The bacteria were also growth in the presence of 50 nM hemin or 20 μM FeCl_3_. The data presented are the mean ± standard deviation of the endpoint OD600nm measured at 24 h. Each data point represents a separate biological replicate from three independent experiments. Statistical significance was determined using an ordinary one-way ANOVA with a tukey’s multiple comparison test. ***p<0.001, ****p<0.0001.

**Figure S3. The *fhuC* gene of *S. lugdunensis* is required for hydroxamate and staphyloferrin siderophore utilization.** Show is the mean diameter of growth for the *fhuC* mutant carrying either vector control or the p*FhuC* plasmid. Growth was measured around sterile paper discs that were impregnated with the indicated iron source or sterile Milli-Q water as a negative control. The data from the plate bioassays demonstrate that a *S. lugdunensis fhuC* mutant can use a catecholamine siderophores but cannot utilize hydroxamate-bound or staphyloferrin-bound iron. The hydroxycarboxylates staphyloferrin A (SA) and staphyloferrin B (SB) were administered as culture supernatants derived from an *S. aureus* Δ*sbn*Δ*sfa* mutant strain unable to produce siderophore as well as *in-vitro* synthesized siderophores. DFO; Desferrioxamine B. Data are the average of at least three independent experiments.

**Figure S4. IsdL-dependent heme acquisition from hemoglobin in *S. lugdunensis*.** In A) the growth of *S. lugdunensis* N920143 wildtype (WT) and ATPase deficient mutants Δ*isdL*, Δ*fhuC and* Δ*isdL*Δ*fhuC* is shown. Strains were grown in the presence of 2.5 µg/mL human hemoglobin (hHb). 500 µL of RPMI + 1% casamino acids + 10 µM EDDHA were inoculated to an OD_600_ of 0.005 in 48 well plates, OD_600_ was measured every 15 min in an Epoch2 plate reader. For reasons of clarity, values after 24 hrs are shown. Mean and SD of three experiments are displayed. Statistical analysis was performed using one-way ANOVA followed by Dunnett‘s test for multiple comparison. B) shows the BACTH assay (Euromedex) of the permease IsdF and the ATPases IsdL and FhuC (*S. lugdunensis* N920143). The proteins were cloned into pKT25 and pUT18C, respectively, and co-transformed into *E. coli* BTH101. As negative control empty vectors (-), as positive control leucine zippers (zip) were used. BTH101 strains were streaked onto LB agar containing X-Gal IPTG ampicillin and kanamycin and incubated for 2 days at 30°C. Blue colour indicates protein-protein interaction.

**Figure S5. Clinal isolates of S. lugdunensis lacking the sst-1 locus fail to utilize norepinephrine to support growth.** The ability of the identified *S. lugdunensis* clincal isolates to utilize norepinephrine (NE) as an iron source is shown. The bacteria were grown in RPMI with 1% (v/v) casamino acids and 0.1% (v/v) heat inactivated horse serum. NE was added at 50 μM and FeCl_3_ was used as a control at 20 μM. The data shown are the mean ± standard deviation of the endpoint optical density at 600nm (OD600nm) measured after 24 hr. The data derive from three independent experiments and each symbol represents a separate biological replicate. Statistical significance was determined by ordinary one-way ANOVA with a tukey’s post-test. In B and D, n.s. indicates not significant, *p<0.05, ****p<0.0001.

**Figure S6. Casamino acids contain factors that are utilized by *S. lugdunensis* and *S. aureus* as iron sources in an Sst-dependent manner**. In (A) growth of wildtype *S. lugdunensis* and the Δ*sst*-1/2 mutant in RPMI supplemented with 1% (w/v) casamino acids in the absence of any other chelator (i.e., horse serum) is shown. The data shown are the mean ± standard deviation of the measured OD600nm after 24h. Bacteria were pre-cultured in serum free RPMI overnight in the presence of antibiotics to maintain plasmid selection. The addition FeCl_3_ was used as a control to show the inability of the Δ*sst*-1/2 mutant to grow is due to Fe starvation. Statistical significance as determine using an ordinary one-way ANOVA with a Dunnett’s multiple comparison test using wildtype as a comparator. In (B) growth of *S. aureus* strain Newman and siderophore biosynthesis/utilization mutants in RPMI with 1% (w/v) casamino acids is shown. Bacteria were pre-cultured in serum free RPMI overnight to deplete the cells of Fe and then inoculated into RPMI + casamino acids in the presence of increasing concentration of horse serum. The data shown are the mean ± standard deviation of the measured OD600nm after 24h. Statistical significance as determine using an ordinary one-way ANOVA with a Dunnett’s multiple comparison test using each wildtype *S. aureus* Newman at a given horse serum concentration as a comparator. In (A) and (B) n.s denotes not significant and ****p<0.0001.

